# Distinct optimization of motor control and learning: Motor learning achieves different motor patterns from overtrained self-paced movements

**DOI:** 10.1101/2024.10.23.619779

**Authors:** Takuji Hayashi, Ken Takiyama

## Abstract

Self-paced movement, refined through extensive training and optimization, serves as an ideal model for investigating internal objectives within the sensorimotor system without external constraints. Previous studies have demonstrated that self-paced movement is consistent within individuals but variable across individuals, emphasizing a robust and unique optimization in self-paced motor control. While motor learning is also an optimization process, it remains unclear whether the internal objectives overlap between motor control and learning, or, more specifically, whether motor patterns identified in self-paced motor control are replicated in motor learning. Given that optimality principles typically address redundancy, human behavioral experiments were designed to equate movement distances for both motor control and learning in self-paced movements and to thus compare redundant motion parameters, movement velocity and duration. We dissected participants’ reaching movements at their preferred pace, controlling a visual cursor to a target on a display over specified movement distances, through a detailed analysis that that accommodates nonlinear interaction and significant inter-individual baseline differences of movement velocity and duration. In self-paced motor control, participants adjusted both velocity and duration in response to changes in target distances. However, when visuomotor shift perturbations were applied to the cursor in the longitudinal direction, triggering motor learning, participants adapted their movement distance solely by modulating velocity, while leaving duration unchanged. These results suggest that distinct optimization strategies are employed in motor control and learning to resolve movement redundancy.

**Significance statement:** Natural behaviors are thought to be well-optimized. Self-paced movements—an important form of such behaviors—showing significant stability within individuals but variability across individuals. This suggests that the sensorimotor system in the central nervous system operates through a unique and robust optimization process for self-paced motor control. Self-paced motor control is further refined through motor learning. However, it remains unclear whether motor control and learning share a common objective function. Our findings demonstrate that laboratory-based motor learning does not replicate the motor patterns observed in overtrained, self-paced movements performed under natural conditions. This suggests that distinct optimization mechanisms are at play in motor control and learning, indicating that these processes are not contiguous.

## INTRODUCTION

In guiding movements, the sensorimotor system in the central nervous system employs visuomotor mapping to convert visual information into motor commands (Hayashi et al., 2016; Schmidt, 1975). For instance, golfers learn to adjust their club-head motion to control ball distance based on the hole’s proximity. Conditions on the green, such as an unexpected, steep slope, may cause the ball to travel farther than intended, prompting adjustments in subsequent putts. The error between the expected and actual ball positions can lead golfers to modify their club-head motion, potentially leading to a standardized motion for both longer putts on steep slopes and shorter ones on flat surfaces. However, it remains uncertain whether motor commands aimed at the visual target and those adapted from visual errors result in identical motor patterns.

The sensorimotor system aims to minimize costs and maximize rewards when executing movements (Todorov, 2004). Self-paced movements, devoid of external constraints, serve as an ideal model for investigating these internal objectives. The principle of optimality often resolves redundancy (Todorov, 2004; Todorov and Jordan, 2002), as reflected in the velocity and duration of movements; Despite constant movement distances, kinematic adjustments in velocity, duration, or both are necessary. Previous studies have demonstrated that both velocity and duration significantly vary with the distance to the movement target (Gordon et al., 1994). Self-paced movements are consistent within but inconsistent across individuals (Berret et al., 2018; Labaune et al., 2020). These studies suggest that a robust and unique motor control optimization is evident in long-term, overtrained self-paced movements targeting different distances. When internal optimization aligns with motor control and learning, the learned motion parameters mirror those generated in self-paced motor control across varying distances.

This study investigates the impact of motor learning to modify movement distance on movement velocity and duration, in comparison to self-paced motor control guided by visual cues. Participants performed self-paced reaching movements, adjusting both velocity and duration individually. The experiment induced sensorimotor adaptation using a visuomotor shift perturbation (Krakauer et al., 2000; Pearson et al., 2010) that altered the visual cursor’s distance, representing hand position, either increasing or decreasing it. Our refined analysis allowed us to minimize the impact of substantial baseline inter-individual differences and nonlinear interactions between velocity and duration, enabling a clearer investigation of these adjustments. Notably, our results revealed that visuomotor perturbations uniquely influence velocity, whereas changes in target distance affect both parameters. These findings suggest that the sensorimotor system employs distinct optimization strategies for motor control and learning, resolving the inherent redundancy.

## RESULTS

### Refined Analysis of Movement Velocity and Duration

Understanding movement velocity and duration in self-paced movements poses significant challenges. One such challenge is redundancy: multiple combinations of velocity and duration can achieve the same movement distance. For instance, compared to a 100-mm reaching movement, a 120-mm movement can be accomplished by increasing velocity, duration, or both by a factor of 1.2. Thus, there are multiple solutions for any given distance. Additionally, if only velocity is modified, a hypothetical baseline of 1 mm/ms requires an increase of 0.2 mm/ms, while a hypothetical baseline of 3 mm/ms requires an increase of 0.6 mm/ms—three times the velocity adjustment. Consequently, the baseline motor patterns, which vary widely across individuals (Berret et al., 2018; Labaune et al., 2020), influence the necessary adjustments. Furthermore, while the relationship between distance, velocity, and duration in constant-speed movements can be captured by a simple equation (distance = velocity × duration), the actual trajectory of reaching movements follows a smooth, bell-shaped profile in the velocity domain (Abend et al., 1982; Morasso, 1981). The height and width of this bell-shaped curve, representing preferred velocity and duration, systematically vary with target distance (Berret et al., 2018; Labaune et al., 2020). Previous theoretical studies that successfully modeled bell-shaped velocity profiles explicitly fixed the movement duration at the outset (Flash and Hogan, 1985; Harris and Wolpert, 1998; Todorov and Jordan, 2002; Uno et al., 1989). However, these models do not explain how movement velocity and duration are adjusted in motor learning. To date, no analysis has fully resolved these issues.

To address these challenges, we developed a three-step analysis. First, we modeled the bell-shaped velocity curve using a Gaussian function: *V*(*t*) = *ν* exp (−(*T*−*t*)^2^/2*τ*^2^), where velocity (*V*) is described as a function of time (*t*), with the timing of peak velocity (*T*). The parameters representing height (*ν*) and width (*τ*) correspond to velocity and duration, respectively. Second, we calculated the movement distance by integrating the Gaussian function, 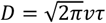 (where *D* is distance), yielding a structure analogous to the constant-speed case, i.e., the product of velocity and duration. Finally, we applied a log transformation to the Gaussian integral: 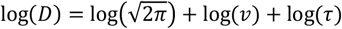. This linearization of the nonlinear interaction allows us to evaluate adjustments in velocity and duration across different motor patterns. Overall, this analysis provides a robust framework for examining velocity and duration in distance-based sensorimotor adaptation, effectively addressing the challenges outlined.

### Changes in Movement Velocity and Duration Across Target Distances

We examined the efficacy of the proposed analysis in self-paced movements across various target distances in Experiment 1. Right-handed participants (n = 24, 13 males and 11 females, ages ranging from 20 to 46) performed forward-reaching movements with their preferred velocity and duration, maintaining a consistent pace throughout the experiment (Fig. 1a and b). The target was located at distances of [60, 70, 80, 90, 100, 110, 120, 130, and 140] mm from the start position, presented in a pseudo-random order (Fig. 1c).

**Figure 1:**
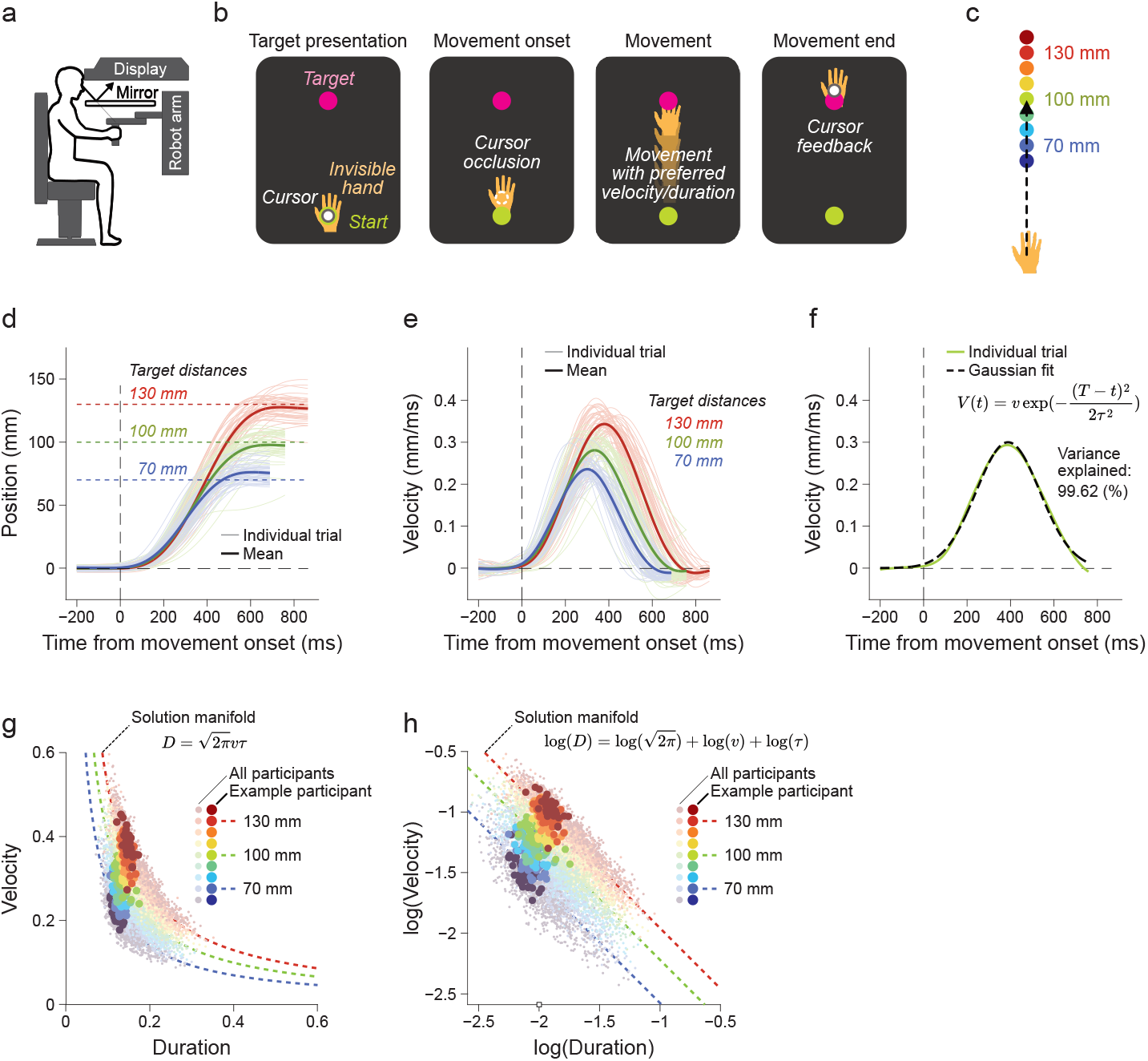
Experimental settings and data analysis. a. Participants performed point-to-point reaching movements while holding the robot manipulandum. b. Verbal instructions encouraged participants to initiate movements with their preferred velocity and duration. Visual feedback was not provided during the movements but was shown at the end. c. The visual target distances were randomly varied trial-by-trial between 60 mm and 140 mm during Experiment 1. d,e. Position (d) and velocity (e) profiles towards 70 mm (blue), 100 mm (green), and 130 mm (red) for a representative participant. Thin light lines indicate individual trials, while the thick dark lines represent the mean. f. The Gaussian fitting (dashed black line) of the example velocity profile showed high accuracy (99.62% variance explained). g. Curved solution manifolds (colored dashed lines) were used to achieve the target distances on the velocity-duration plane of the Gaussian fitting. Each individual trial (dots representing all participants; solid dots for the representative participant from d and e) lies on the manifolds. h. Log transformation, which linearizes the manifolds, allows for a fair evaluation of motion parameters despite large inter-individual differences. From the scatter plot, we computed the preferences and slopes for each participant (Fig. 2).

Participants aimed to align the visual cursor, representing hand position, with these targets, despite the cursor being occluded during the movements. This procedure was tested in three phases: the beginning, middle, and end of the experiment, interspersed with other motor tasks, including the sensorimotor adaptation task shown below. Figure 1d and e show the time course of endpoint position and velocity for three target distances, [70, 100, and 130] mm, for a representative participant. The endpoints exhibited consistent, smooth trajectories without online visual feedback (Fig. 1d). The velocities followed a bell-shaped pattern and scaled with increasing target distances (Fig. 1e). The trajectory scaling suggests that both movement velocity and duration varied with target distance.

Utilizing the Gaussian function with the bell-shaped curves resulted in significant fitting accuracy, with 97.8 ± 0.2% (mean ± SEM) of the variance explained (Figs. 1f and S1a). Notably, the predicted distances from the Gaussian integral were consistent with the measured distances (slope: 0.9812 ± 0.0034, intercept: −0.0012 ± 0.0004, Fig. S1b). For any given target distance, there are multiple possible combinations of movement velocity and duration. The data points from all participants were scattered across the solution manifolds (Figs. 1g and h). The raw fitting parameters, *ν* and *τ*, represent nonlinear interactions (Fig. 1g), whereas applying a log transformation to *ν* and *τ* makes these interactions linear (Fig. 1h). The log transformation linearizes the relationship, allowing for the evaluation of velocity and duration, as it resolves the nonlinear dependency and ensures that the weights of velocity and duration are equal on this plane (i.e., 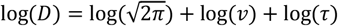. Therefore, we used log(*ν*) and log(*τ*) as measures of subjective preferred velocity and duration.

We found that, in the self-paced reaching movements, the movement distances varied with target distances (Fig. 2a). However, the resultant movement distances were slightly overshot for the 70-mm target (73.7 ± 0.1 mm) and undershot for the 130-mm target (129.8 ± 0.1 mm). These small shifts toward the mean of the overall movements may have been induced by movement history (Hammerbeck et al., 2014), where the subsequent movement was adjusted to resemble the previous one. Specifically, since the target distances just prior to the 70-mm target were often larger, with theoretical average of 103.75 mm, than 70 mm, movements toward the 70-mm target became slightly farther. We also found that the log velocity (Fig. 2b) and log duration (Fig. 2c) changed with target distance. The log velocity increased by 0.40 ± 0.01 from the 70-mm to the 130-mm target distance, while the log duration increased by 0.16 ± 0.01, indicating that movement velocity was more heavily weighted than duration on the log plane (t(23) = 15.07, p = 2.08 × 10^−13^). To examine the stability of self-paced movements within individuals, we computed two representative values: preference, indicating which velocity and duration were preferred (Fig. 2d), and slope, indicating the ratio of how much velocity and duration changed (Fig. 2e, see Methods). We found that preference was significantly biased toward velocity (Fig. 2d, t(23) = 6.77, p = 6.60 × 10^−7^) and was highly consistent within individuals (Fig. 2f, intra-individual standard deviation vs. inter-individual standard deviation: 0.14 ± 0.02 vs. 0.38, t(23) = 14.03, p = 9.22 × 10^−13^). The slope was consistently greater than 45 degrees for all individuals (Fig. 2e, t(23) = 20.05, p = 4.58 × 10^−16^) and was also highly consistent within individuals (Fig. 2g, intra-individual standard deviation vs. inter-individual standard deviation: 3.90 ± 0.53 vs. 5.25, t(23) = 2.53, p = 0.019). There was no correlation between preference and slope (Fig. 2h, r = 0.12, p = 0.57). Participants who preferred faster movement did not necessarily adjust their velocity, suggesting that individual preferences for velocity and duration are independent of how the two are adjusted based on target distances. These results demonstrate that our analysis effectively captured self-paced movements, and that movement velocity and duration were consistent within individuals but varied across individuals.

**Figure 2:**
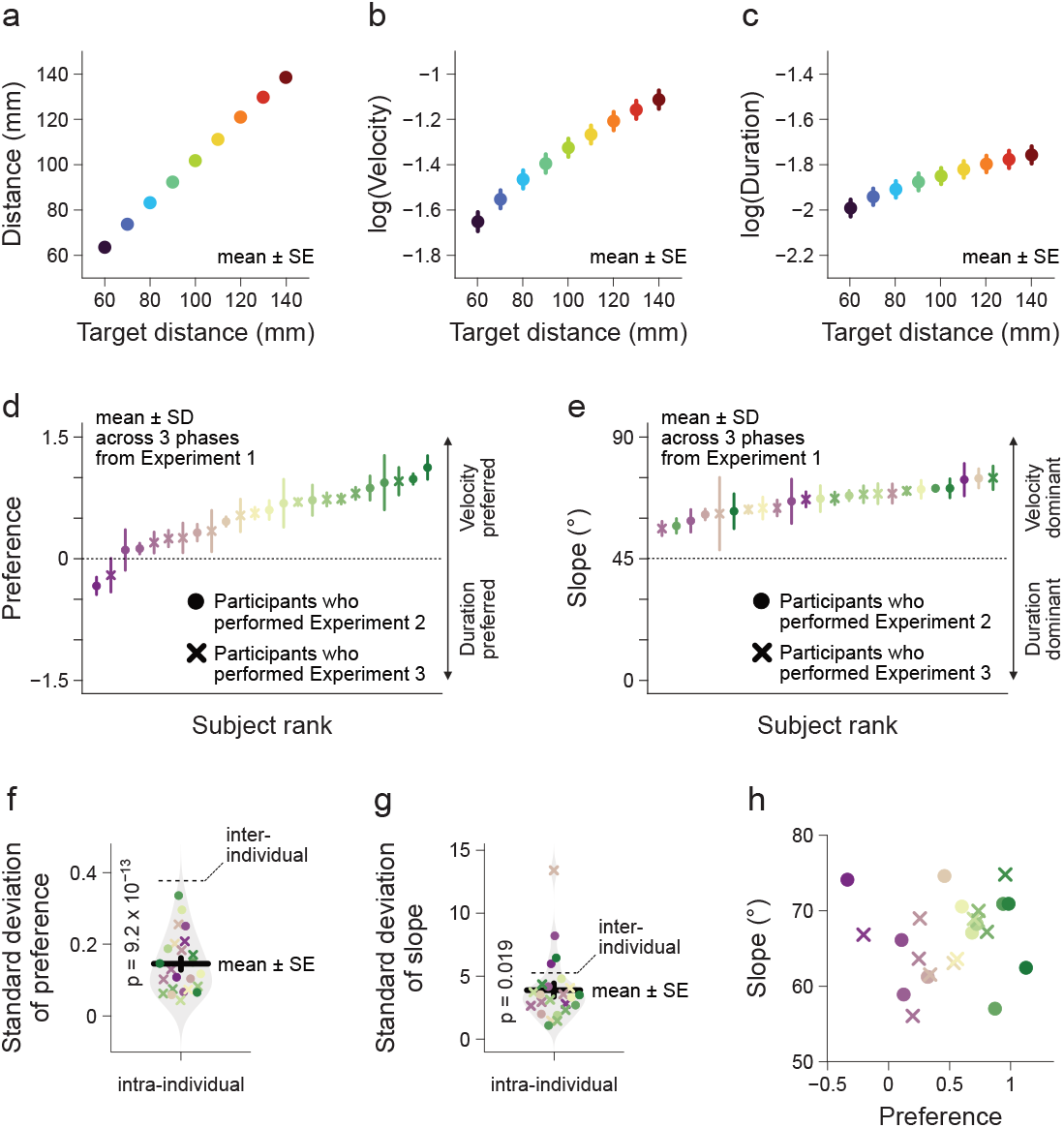
Target position determines movement velocity and duration. a-c. The nine target distances influenced movement distances (a), velocities (b), and durations (c). d,e. We computed two indices: preference (d), which is the difference between mean movement velocity and duration, and slope (e), which represents the angle of the black line as shown in Figure 1h. Colors indicate participant rank, sorted by preference (d), and are also used for the slope (e). f,g. Intra-individual standard deviation, represented as error bars in d and e, was smaller than the inter-individual standard deviation for both preference (f) and slope (g). h. The preference was not correlated with the slope (color coding is the same as in d).

### Not Movement Duration but Movement Velocity Changes in Sensorimotor Adaptation

In Experiment 2, we investigated how the sensorimotor system resolves the redundancy between movement velocity and duration during sensorimotor adaptation. A 30-mm shift-up (Fig. 3a) or 30-mm shift-down perturbation (Fig. 3b) was introduced to 12 participants (seven males and five females, aged between 21 and 37). The target was consistently located at 100 mm, so ideal learning would result in movement distances of 70 mm (down modulation) and 130 mm (up modulation). Participants performed 200 learning trials, with either visuomotor shifts before or after the null trials (comprising 20 baseline trials and 50 washout trials) (Fig. 3c). Each participant experienced both shift-up and shift-down perturbations in a counterbalanced order across Experiment 1.

**Figure 3:**
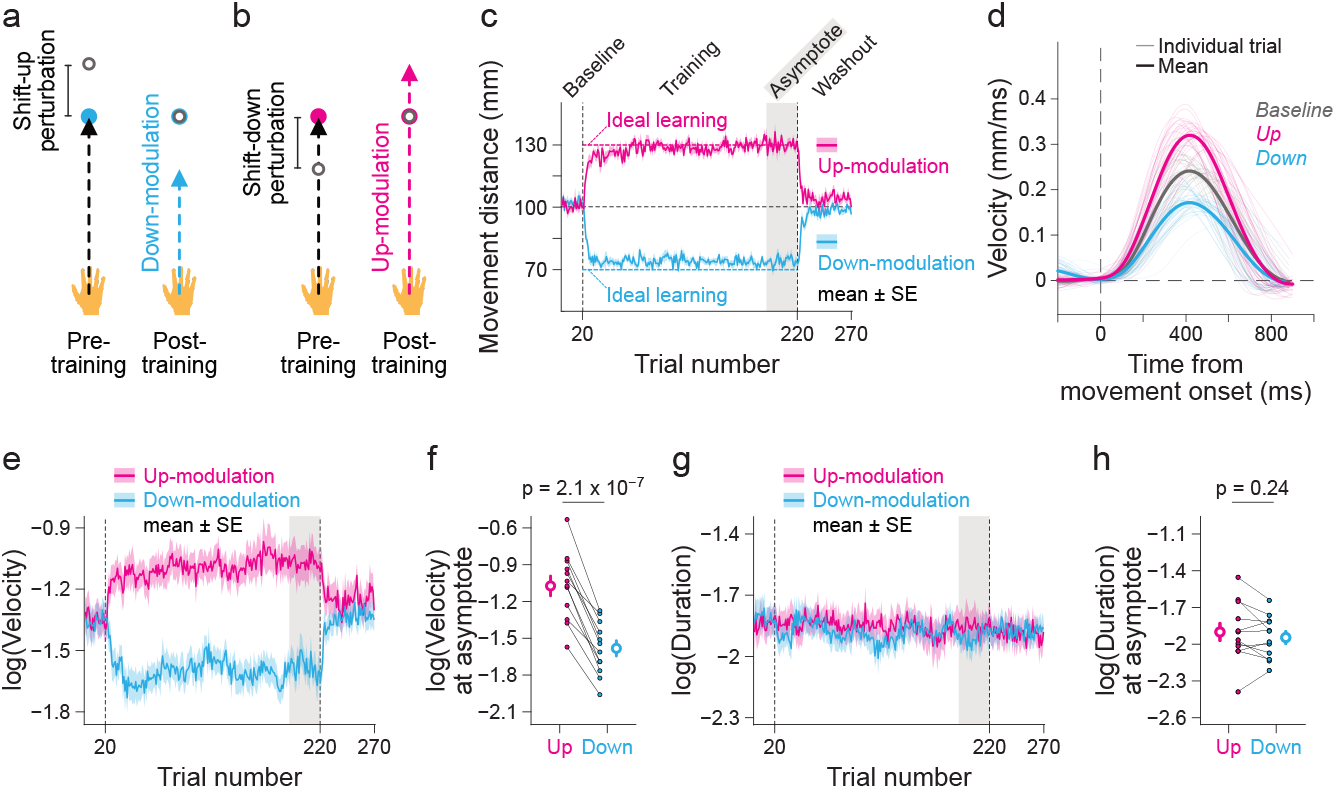
Visuomotor shift modulates only movement velocity. a,b. Visuomotor shift perturbations were introduced in the longitudinal direction, modifying movement distances to be shorter (a, down-modulation) or farther (b, up-modulation). c. The visuomotor shifts correctly adjusted movement distances (thick lines) to the ideal levels (thin broken lines). d. In the baseline (gray) and resultant velocity profiles of a representative participant (magenta and cyan), the height (velocity) appears to change, while the width (duration) remains unchanged. e,f. Movement velocity was modulated in response to the movement distance adjustment. g,h. In contrast, no modulation of movement duration was observed.

In sensorimotor adaptation, perfect learning in the distance domain does not occur (Al-Fawakhiri and McDougle, 2024). During the final 30 trials of the learning phase, movement deviations averaged 74.1 ± 0.8 mm for down-modulation and 129.9 ± 0.8 mm for up-modulation (Fig. 3c). These results were not statistically different from the movements toward the 70-mm and 130-mm targets in Experiment 1 (Fig. 2a, interaction of [Experiment 1 vs. Experiment 2] × [70-vs. 130-mm target distances], F(1,11) = 0.09, p = 0.78). Log movement velocity adjusted significantly due to sensorimotor adaptation (Fig. 3d->f; −1.58 ± 0.06 for down-modulation, −1.07 ± 0.08 for up-modulation, t(11) = 11.33, p = 2.10 × 10^−7^). In contrast, log movement duration did not differ significantly between down-modulation (−1.90 ± 0.05) and up-modulation (−1.85 ± 0.08) (Fig. 3d, g, h, t(11) = 1.25, p = 0.24). These results indicate that sensorimotor adaptation to visuomotor shifts primarily affects velocity rather than duration, highlighting distinct optimization strategies in motor control and learning.

### Changes in Target Distances to Match Movement Volatility

We found that both movement velocity and duration changed with the target distance in Experiment 1(Fig. 2b and c), whereas only velocity changed with sensorimotor adaptation in Experiment 2 (Fig. 3e-h). Beyond motor control and learning, an additional methodological difference concerns the volatility of movement distances. In Experiment 1, target distances varied on every trial, leading to corresponding changes in movement distances. In contrast, during sensorimotor adaptation in Experiment 2, movement distances remained constant despite the perturbations.

To test the possibility that movement volatility causes an imbalance in velocity and duration adjustments, we used the mean movement distances from Experiment 2 as the target sequence for Experiment 3 with the remaining 12 participants (Fig. 4a; six males and six females, aged between 20 and 46). This design allowed us to investigate whether target distances alter both movement velocity and duration under the same movement volatility observed in sensorimotor adaptation. The movement distances followed the target sequences precisely (Fig. 4b, 76.1 ± 0.9 for down-modulation, 131.8 ± 1.0 for up-modulation, t(11) = 96.95, p = 1.76 × 10^−17^). Additionally, we observed changes in both log movement velocity (Fig. 4c-e, −1.58 ± 0.06 for down-modulation, −1.12 ± 0.06 for up-modulation, t(11) = 23.41, p = 9.82 × 10^−11^) and log movement duration (Fig. 4c, f, g, −1.88 ± 0.05 for down-modulation, −1.79 ± 0.05 for up-modulation, t(11) = 4.71, p = 6.38 × 10^−4^). The results demonstrate that movement volatility, driven by target distances, does not disrupt the balance between adjustments in movement velocity and duration. This highlights the distinct optimization processes in motor control and learning.

**Figure 4:**
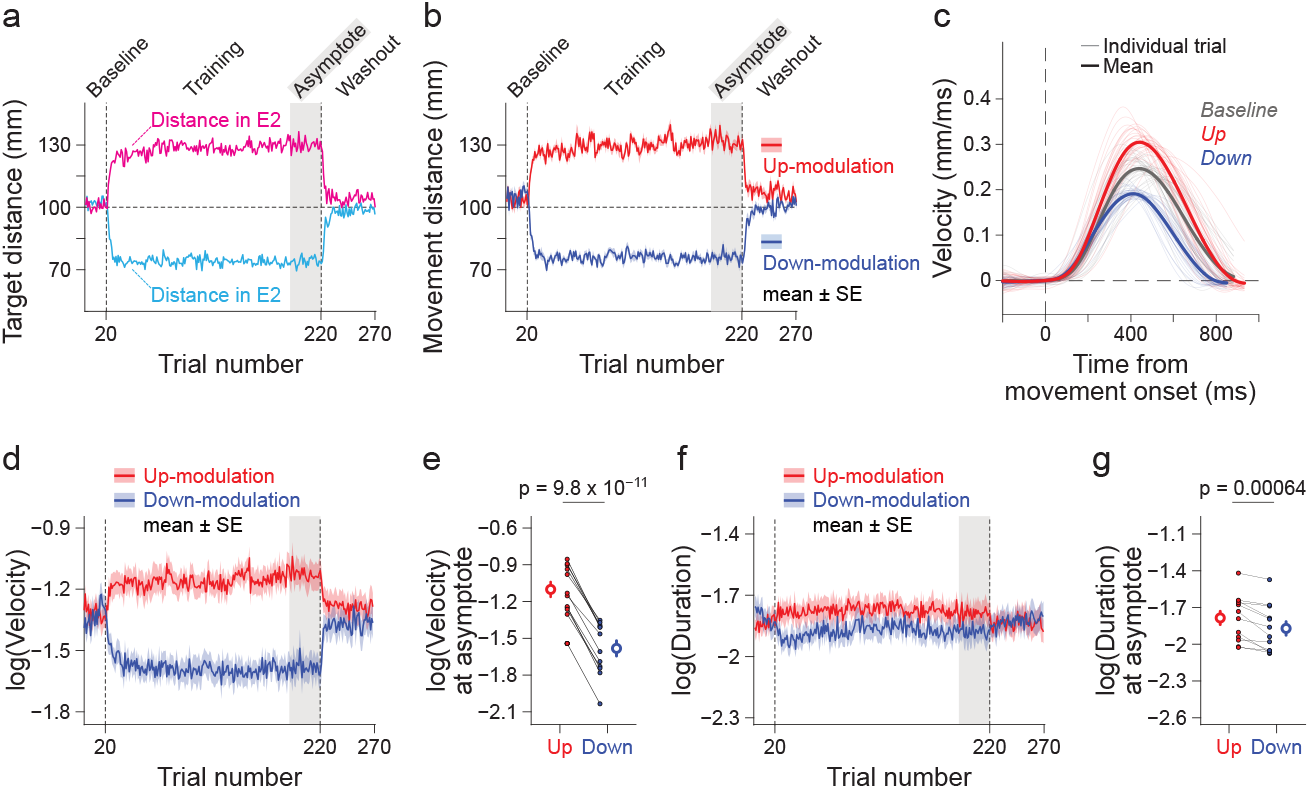
Target position with matched movement volatility modulates both velocity and duration. a. To match movement volatility to sensorimotor adaptation, we used the mean movement distances observed in Experiment 2 (E1) as the target distances in Experiment 3. b. Movement distances clearly changed, similar to those observed during sensorimotor adaptation. c. In the baseline (gray) and resultant velocity profiles of a representative participant (red and blue), both the height (velocity) and width (duration) appeared to change. d-g. In contrast to sensorimotor adaptation, both movement velocity (d, e) and duration (f, g) changed. These panels are analogous to those in Figure 3.

## DISCUSSIONS

The sensorimotor system generates natural behavior autonomously without external constraints, which serves as a fundamental model to elucidate the internal objective function. This optimality principle often helps resolve the redundancy inherent in the sensorimotor system. Interestingly, both movement velocity and duration, though redundant in a self-paced movement, exhibit consistent patterns within individuals while varying across individuals (Fig. 2) (Berret et al., 2018; Labaune et al., 2020). This suggests that individuals follow unique, robust optimization strategies. Self-paced motor control is refined through motor learning. However, it remains unclear whether motor learning shares the same optimization principles as motor control. By comparing redundant movement parameters, resolved through the optimization process, we investigated whether motor learning replicates the motor patterns identified in self-paced motor control. When internal optimization aligns with both motor control and learning, the learned motion parameters should mirror those generated during motor control across varying distances.

The human behavioral experiment was designed to assess two redundant motion parameters in motor control and learning. Participants performed self-paced reaching movements toward targets set at 100-mm distances and adapted to 30-mm shifts, either up or down, which could be corrected by changes in movement velocity, duration, or both. Our results revealed that motor learning, induced by visuomotor shift perturbations, exclusively altered movement velocity (Fig. 3). In contrast, motor control adjustments for different target distances affected both velocity and duration (Figs. 2 and 4). This discrepancy in motion parameters between motor control and learning suggests that each is governed by distinct optimization processes.

Previous studies often combined movement velocity and duration into a single index, termed movement vigor (Shadmehr and Ahmed, 2020), without distinguishing between the two. Various factors have been proposed to determine movement vigor, including expectations of reward (Kawagoe et al., 1998; Summerside et al., 2018; Takikawa et al., 2002), metabolic energy expenditure (Shadmehr et al., 2016; Wong et al., 2021), mechanical smoothness (Flash and Hogan, 1985; Todorov and Jordan, 1998; Uno et al., 1989), sensorimotor noise (Harris and Wolpert, 1998), and environmental uncertainty (Burdet et al., 2001). However, these studies typically constrained movement vigor by explicitly setting either movement velocity or duration, overlooking biologically natural, self-paced behavior and making it difficult to observe internal objectives directly. Notably, Huh and Sejnowski (2016) proposed a potential internal objective for self-paced movement, called “drive,” which refers to the motivational level that governs movement vigor throughout trajectory curvatures and sizes (Huh and Sejnowski, 2016). This concept aligns with our findings, which demonstrate low intra-individual variability in preferences and slopes during self-paced motor control (Fig. 2d-g). Future research should seek to identify the unique objectives driving motor learning.

The “as quickly as possible” instruction has been widely used to investigate human motor control and its computational models (Fitts and Peterson, 1964; Gordon et al., 1994; Kelso et al., 1979; Meyer et al., 1988; Reppert et al., 2018; Wong et al., 2021), as has the “comfortable-speed” instruction (Abend et al., 1982; Berret et al., 2018; Fitts, 1954; Huh and Sejnowski, 2016, 2015; Labaune et al., 2020; Morasso, 1981; Todorov and Jordan, 1998; Wang et al., 2016; Young et al., 2009). Al-Fawakhiri and McDougle (2024) found that sensorimotor adaptation alters both movement velocity and duration under the “as quickly as possible” instruction (Al-Fawakhiri and McDougle, 2024), which contrasts with our findings, where no change in movement duration was observed under the comfortable-speed instruction. This suggests that external constraints on movement vigor may significantly influence motion parameters, potentially overriding internal optimization objectives.

Motor commands are modulated by visuomotor mismatches, as demonstrated by numerous studies (Danziger and Mussa-Ivaldi, 2012; Franklin and Wolpert, 2008; Wolpert et al., 1994). Although visual distortion at the movement midpoint does not affect the success of target acquisition, it prompts corrective adjustments in hand motion to realign with visual errors, suggesting that visual information takes precedence over motor and proprioceptive feedback. One computational theory proposes that movement duration is determined by the task’s accuracy or success rate at movement completion (Tanaka et al., 2006). Our results support this theory, showing that while movement velocity is flexible, movement duration remains fixed to the visual target distances. This underscores that the sensorimotor system optimizes internal objectives in motor planning based on visual information during both motor control and learning. However, even if this explanation holds, a key question arises: If visual information determines these redundant motor control patterns, such as velocity and duration, how and why do the individual patterns become crystallized through motor learning? Further exploration is required to address this mystery.

## MATERIAL and METHODS

### Participants

Twenty-four right-handed participants (13 males, 11 females, aged 20–46 years) took part in the experiments. All participants completed Experiment 1. Of these, half (7 males, 7 females, aged 21–37 years) performed Experiment 2, while the other half (6 males, 6 females, aged 20–46 years) completed Experiment 3. Both Experiments 2 and 3 were interleaved with Experiment 1. Informed consent was obtained from all participants, and the study was approved by the ethical committees of the University of Tokyo.

### General Task Settings

Participants performed point-to-point reaching movements while holding a robot manipulandum (KINARM) with their right hand. Seated in front of a monitor and facing it via a mirror angled upwards, they viewed the screen through the mirror while performing movements with their hands beneath it (Fig. 1a). This setup occluded direct visual access to their hands, reducing awareness of the visual-motor mismatch. A visual target appeared 200 ms after the visual cursor, which represented the hand position, reached the start position. The cursor then disappeared, prompting participants to perform the reaching movement without online visual information (Fig. 1b). The movement ended when the threshold of <0.03 mm/ms was met for 250 ms, after which the cursor reappeared either with or without visuomotor shift perturbations. During backward movements, driven by forces from the KINARM, the cursor remained visible at the movement endpoint without additional feedback from the participants’ hand movements. Handle positions were recorded at a 1000 Hz sampling rate.

In the experiment, participants completed three consecutive sessions of Experiment 1, interspersed with either ‘up-modulation’ or ‘down-modulation’ sessions from Experiment 2 or 3. For example, the sequence consisted of Experiment 1, up-modulation from Experiment 2, Experiment 1, down-modulation from Experiment 2, and then a final Experiment 1 phase. A one-minute break was given every 5 minutes, during which participants completed an average of 137.9 ± 4.2 trials (mean ± SEM). Participants were instructed to perform movements at their preferred velocity and duration, focusing on accuracy without deliberately adjusting their pace. This ensured that movements remained consistent within each individual but varied between individuals throughout the experiment (Fig. 2).

### Experiment 1: Self-paced Movements Toward a Target with Trial-by-Trial Distance Changes

In Experiment 1, the target location varied trial-by-trial across nine possible distances—[60, 70, 80, 90, 100, 110, 120, 130, and 140] mm—presented in pseudo-random order (Fig. 1c). Participants completed a block of nine trials 15 times, for a total of 135 trials in each of the three phases. Prior to the experiment, participants practiced these movement distances.

### Experiment 2: Movement Distances Modulated by Visuomotor Shift Perturbation

Twelve participants performed adaptation tasks. The target remained fixed at 100 mm throughout the experiment. Initially, no visual shift occurred; however, at the end of each movement, the visual cursor shifted 30 mm forward or backward along the longitudinal axis relative to the actual hand position. This shift resulted in an ideal movement distance of either 70 mm (down-modulation) or 130 mm (up-modulation). Each participant experienced both types of visuomotor shift perturbations, with the sequence counterbalanced. Participants completed 20 trials without shifts (baseline), followed by 200 trials with perturbations, and concluded with 50 trials without shifts (washout).

### Experiment 3: Movement Distances Modulated by Adaptive Target Shift

The other twelve participants performed tasks with different target sequences, without the visuomotor shift perturbations. To match the movement volatility observed in Experiment 2, the mean movement distances from Experiment 2 were used as the target distances in Experiment 3. Additionally, the order of up- and down-modulation was counterbalanced across these participants.

### Data Analysis

For movement distances, we used the location at movement completion. To independently estimate movement velocity and duration, we fitted a Gaussian function to the velocity profiles over time (Fig. 1f). The Gaussian function provided estimates for movement velocity (*ν*) and duration (*τ*). We applied the Gaussian fitting to the velocity profiles from cue presentation to target acquisition in each trial using the Matlab function “fminsearch,” minimizing the mean squared error. To validate this analysis, we compared the measured and predicted movement distances from the Gaussian integral and assessed fitting accuracy using the explained variance (R^2^). Trials with poor fitting accuracy (R^2^ < 90%) were excluded, resulting in the removal of 15.1 ± 1.6 trials (1.6 ± 0.2% of the total) (Fig. S1).

For Experiment 1, we rejected trials as outliers if the movement distance, velocity, or duration deviated from the threshold defined by the median ± 3 interquartile ranges (IQRs) for each target distance within participants. We then computed the representative values, including preference and slope. To determine the preference, indicating which velocity and duration were favored, we calculated the differences in mean velocity and duration in logarithmic space. To assess the slope, which reflects how these parameters change with different target distances, we compared the changes in mean velocity and mean duration across target distances. These calculations were performed separately for the beginning, middle, and end phases in logarithmic space. We then compared the standard deviations within participants (intra-individual variance) and across participants (inter-individual variance) to examine the consistency of motor patterns within individuals. Additionally, we computed the correlation coefficients between preference and slope to explore whether individuals who favored faster movements also adjusted their velocity in response to variations in target distances.

For Experiments 2 and 3, we discarded outlier data points across participants using the threshold defined by the median ± 3 interquartile ranges (IQRs). Additionally, we removed outlier data points within participants from the last 30 trials of the learning phases and computed the mean to evaluate the asymptotic motor patterns of sensorimotor adaptation. Hypothesis testing was performed between the up- and down-modulation conditions or across the Experiments 1 and 2 using a paired sample t-test or a two-way repeated measures ANOVA. Statistical significance was set at p < 0.05.

## Acknowledgments

We thank On Kobayashi and Asako Munakata for helping to organize and run experiments. We thank Daichi Nozaki for the helpful discussions. This work was supported by JSPS KAKENHI grants (JP18K17893, JP23H03296) and JST/PRESTO (JPMJPR23S8) to TH.

## Data availability

The data and codes generated in MATLAB for the current study will be made available upon publication through Zenodo.

**Supplementary Figure 1:**
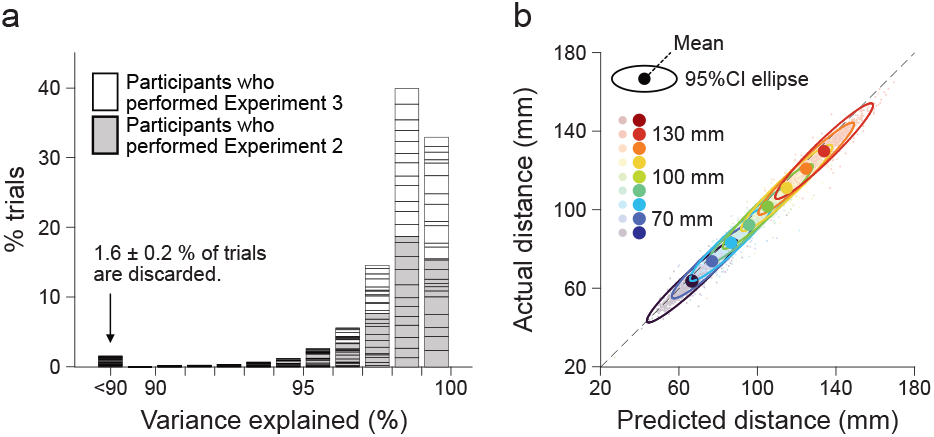
Fitting quality and accuracy. a. For almost all trials, the variance explained was greater than 90%. A small fraction of trials (<90% variance explained) were discarded as outliers (1.6 ± 0.2%). b. The actual movement distances from the data and the predicted distances from the Gaussian integral are highly correlated, as shown by the unity line (black dashed line).

